# Measurement of infection efficiency of a major wheat pathogen using time-resolved imaging of disease progress

**DOI:** 10.1101/318204

**Authors:** P. Karisto, S. Dora, A. Mikaberidze

## Abstract

Infection efficiency is a key epidemiological parameter that determines the proportion of pathogen spores able to infect and cause lesions once they have landed on a susceptible plant tissue. In this study, we present an improved method to measure infection efficiency of *Zymoseptoria tritici* using a replicated greenhouse experiment. *Z. tritici* is a fungal pathogen that infects wheat leaves and causes Septoria tritici blotch (STB), a major disease of wheat worldwide.

We devised an original experimental setup, where we (i) attached living wheat leaves to metal plates allowing for time-resolved imaging of disease progress in planta. Since lesions were continuously appearing, expanding and merging during the period of up to three weeks, daily measurements were necessary for accurate counting of lesions. We also (ii) used reference membranes to characterize the density and the spatial distribution of inoculated spores on leaf surfaces. In this way, we captured the relationship between the number of lesions and the number of viable spores deposited on the leaves and estimated the infection efficiency of about 4 % from the slope of this relationship.

Our study provides a proof of principle for an accurate and reliable measurement of infection efficiency of *Z. tritici*. The method opens opportunities for determining the genetic basis of the component of quantitative resistance that suppresses infection efficiency. This knowledge would improve breeding for quantitative resistance against STB, a control measure considered more durable than deployment of major resistance genes.

## Introduction

Foliar fungal pathogens of plants produce immense numbers of spores (Sache & de Vallavieille-Pope, 1995), but only a small fraction of them causes new infections. Most of the spores do not land on plant tissue that they can infect and are “lost” in the environment. Even among the spores that succeed in finding suitable host tissue, only a moderate proportion will infect and cause lesions. This proportion is determined by the infection efficiency, defined as the probability of an individual spore that lands on a susceptible host tissue to cause a lesion. Infection efficiency is one of the key determinants of the pathogen’s rate of transmission. From the perspective of the host, the host’s ability to reduce the infection efficiency is a major component of quantitative resistance that reduces the rate of epidemic development (Parlevliet, 1979).

In this study, we present a novel method to measure infection efficiency of *Zymoseptoria tritici* (formerly known as *Mycosphaerella graminicola*). This fungal pathogen causes Septoria tritici blotch (STB), a major disease of wheat (*Triticum aestivum*) in Europe (Jørgensen *et al*., 2014) and worldwide (Orton *et al*., 2011). Even controlled epidemics of the disease can lead to notable yield losses if the environmental conditions favour the development of the disease (Fones & Gurr, 2015). STB is controlled mainly by foliar fungicide applications and deployment of disease resistant wheat varieties. However, acquired fungicide resistances are spreading in the genetically diverse population of *Z. tritici* and diminishing the efficacy of fungicide treatments (Fraaije *et al*., 2005; Zhan *et al*., 2006). Similarly, the pathogen has the capacity to adapt to both major gene resistance (Cowger *et al*., 2000) and quantitative resistance (Cowger & Mundt, 2002). Quantitative resistance is generally thought to be more durable than major gene resistance (St.Clair, 2010). Accurate phenotypic characterization of infection efficiency has the potential to improve breeding for quantitative resistance against STB.

The infection cycle of *Z. tritici* starts with the deposition of inoculum composed of wind-dispersed ascospores and rain-splash dispersed pycnidiospores on the leaves (Suffert *et al*., 2011). On the leaf surface, germinating spores penetrate the plant tissue through stomata, enter the apoplast and start to colonise the leaf in an asymptomatic manner. During a latent period of roughly two weeks (Kema *et al*., 1996), the fungus evades recognition by the plant defence mechanisms. Once the fungus has colonised the plant tissue, it starts to produce necrotic lesions within which it produces pycnidia and eventually releases new pycnidiospores that disperse further and generate secondary infections (Orton *et al*., 2011).

Infection efficiency can be quantified as the ratio between the number of lesions visible on a leaf and the number of viable spores deposited on a leaf surface. Therefore, to measure infection efficiency one needs to measure accurately the number of viable spores deposited on the leaf and the resulting number of lesions formed on the leaf. Measurements of infection efficiency were conducted in many airborne fungal pathogens including several rust species. Sache & de Vallavieille-Pope (1995) discuss data on 22 species. Methods used to estimate the number of deposited spores include counting spores directly on the leaf surface (Mehta & Zadoks, 1970; Melching *et al*., 1988), weighing the inoculum (Sache, 1997), and using a known volume of liquid inoculum in which the concentration of spores was measured (Levy, 1989; Shine & Jarriel, 1990).

To achieve a precise measurement of infection efficiency, not only the amount of inoculum but also its spatial distribution across the leaf surface needs to be characterized. When fungal spores arrive to the surface of the leaf in groups, for example as droplets containing several spores, an infection event cannot be with certainty attributed to a single spore. The same argument holds in the case of high densities of spores on the leaf surface when many spores are likely to be present within the area covered by a typical lesion. In these cases, the infection efficiency cannot be determined by simply dividing the number of lesions by the number of deposited viable spores. Moreover, possible interaction between spores when producing lesions may bias the estimate of infection efficiency. Thus, a spatially uniform, precisely measured and sufficiently low density of spores is needed for a reliable measurement. Different approaches to achieve suitable distribution of inoculum include spore settling towers (Brown & Kochman, 1973), atomizers (Statler & McVey, 1987) and paintbrushes (Melching *et al*., 1988).

One of the challenges in estimating the infection efficiency of *Z. tritici* is that spores of this pathogen are not easily visualised on wheat leaves and cannot be counted directly on the leaf surface. The only measurement of infection efficiency of *Z. tritici* available to date was reported by Fones *et al*. (2015). They applied spore suspensions to leaves and spread the inoculum across the leaf manually with a finger covered by a rubber glove. The estimate of the infection efficiency was then given by the ratio between the number of individual lesions observed on the leaves and the total number of spores contained in the inoculated suspension. However, Fones *et al*. (2015) characterized neither the spatial distribution of the deposited spores across the leaf surface nor the viability of spores. In addition, the number of lesions was only measured at a single time point, which may have led to an underestimation of lesion numbers, if not all lesions appeared or if some lesions already merged by this time.

In this study, we developed a method to measure the infection efficiency of *Z. tritici* accurately, with a relatively low effort and low cost. One of the key aspects of the method was the use of reference membranes that provided information on the density and the spatial distribution of viable spores deposited on leaves. In our setup, the leaves were attached to metal plates, so that the inoculated area of the leaves was easy to observe and image during the infection. This allowed us to record the appearance of individual lesions with temporal resolution from the time when first lesions started to appear, which improved crucially the accuracy in counting lesions.

## Materials and Methods

### Plant and fungal material

We planted winter wheat (*Triticum aestivum*) cultivar Drifter seeds in 6×6×11 cm pots containing soil substrate (Jiffy soil substrate GO PP7, Netherlands) and watered them regularly. Cultivar Drifter was classified as susceptible to STB according to multi-year field trials in Germany (Risser, 2010) and also was found to be one of the most susceptible among 335 elite European wheat cultivars exposed to the diverse natural *Z. tritici* population in a recent field experiment in Switzerland (Karisto *et al*., 2018). The plants were fertilized ten days after sowing with 1 l of fertilizer solution (Wuxal Universal-Dünger, Maag-Garden, Switzerland; 1 ml/l diluted in tap water) per 20-24 pots. The plants were grown in the greenhouse with the light/dark cycle of 16/8 hours, the relative humidity of 70 % and the temperature of 18/15 °C. After inoculation, the plants were trimmed twice a week by cutting the newly emerged leaves above the inoculated second leaf.

To prepare fungal spores for plant inoculation, *Z. tritici* strain ST_CH3_99_3D7 (short identifier 3D7; Zhan *et al*., 2002; Septoria tritici blotch network, 2017) blastospores were grown in 50 ml yeast-sucrose-broth (10 g/l sucrose, 10 g/l yeast extract, 50 mg/l kanamycin) for 5 days at 18 °C in the dark. We chose this strain because it is known to be highly virulent on cultivar Drifter under greenhouse conditions (for example Palma-Guerrero *et al*., 2016) and is well characterized both genetically and phenotypically. The liquid culture was then filtered, pelleted and suspended in sterile water and the concentration of spores was determined using a KOVA Glasstic Slide counting chamber. The inoculum was diluted to achieve the required concentrations and 1 ml/l of Tween20 (Biochemica, Applichem Gmbh, Germany) was added. The spore suspension was kept on ice until plant inoculations were conducted on the same day.

### Experimental procedures

The whole experiment was repeated twice as described below. Each repetition can be considered as an independent biological replicate. For clarity, we refer to the two biological replicates as the “first experiment” and the “second experiment”.

We inoculated second leaves of sixteen days old wheat seedlings. For this purpose, the pots were placed into a tray with an aluminium plate in the middle as shown in Fig. 1. The leaves to be inoculated were arranged in four sets containing 5-8 leaves each on an aluminium plate and attached with eight elastic threads. Reference membranes (Whatman™ 3MM Chr Chromatography paper, Fisher scientific) were attached to plates next to each leaf set. Spore suspension was applied while moving the paint gun sprayer (RevolutionAir, Fini Nuair, Italy) twice along the plate. The sprayer was operated with 20 psi pressure (adjusted with the pressure reducer Filterdruckminderer R 1/4”, Einhell, Germany) and the minimal flow rate to maximize the atomization of the spore suspension and the uniformity of coverage.

**Figure 1.**
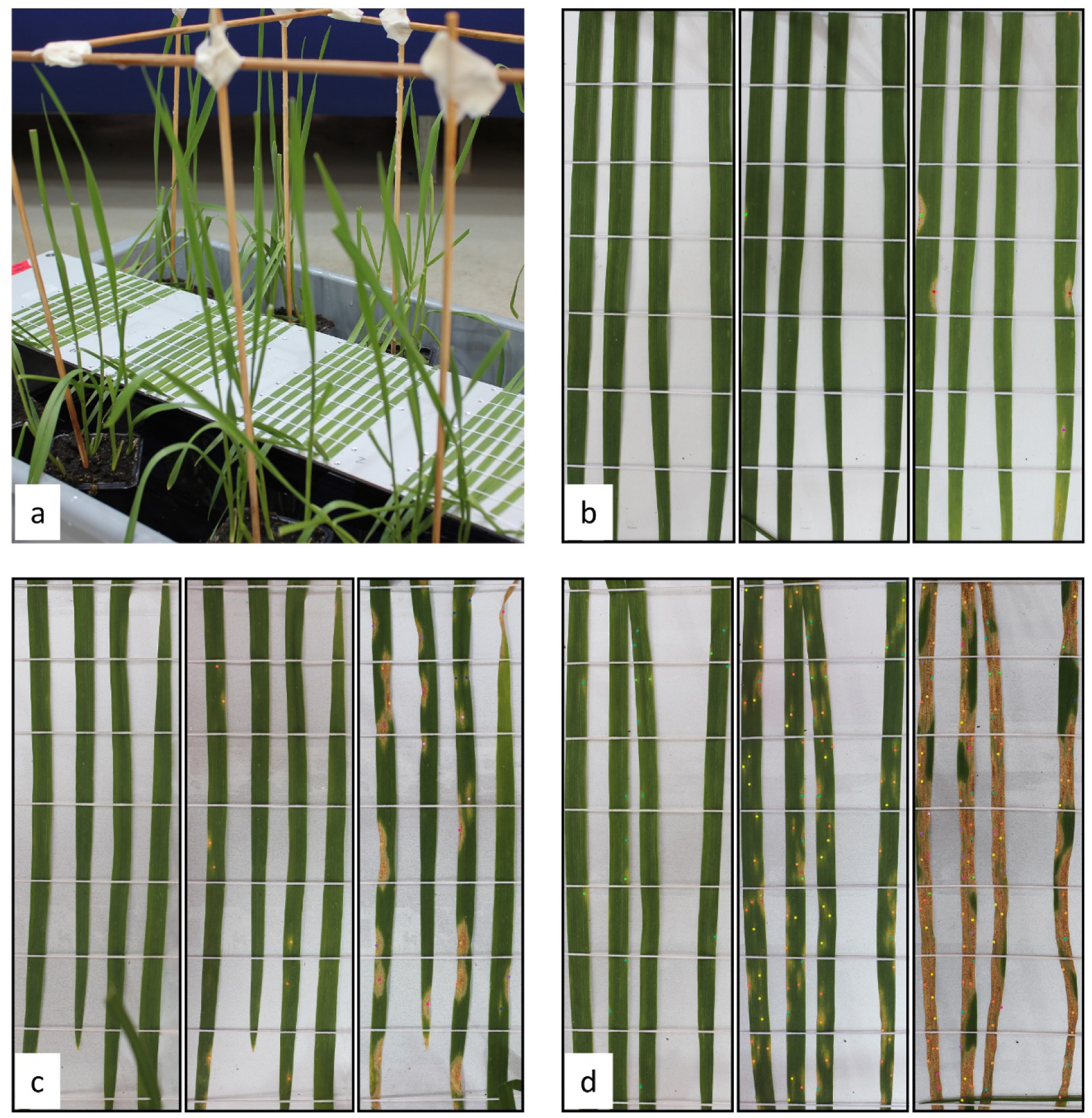
(a) Experimental setup. Second leaves were attached to metal plates by elastic threads. Wooden sticks were used to stabilize individual pots and to support the plastic bags during the high humidity period after the inoculation. Reference membranes were placed on both sides of each leaf set during the inoculation. (b,c,d) Representative leaf sets at 14, 17 and 23 days post inoculation from left to right corresponding to the concentration of spores of 5×10^2^ spores/ml (b), 10^4^ spores/ml (c), and 5×10^4^ spores/ml (d). Lesions are marked with coloured dots that indicate the day and the position of their first appearance.

The plants were infected with spore suspensions that contained 5×10^2^, 10^3^, 5×10^3^, 10^4^, 5×10^4^, and 10^5^ spores/ml. Different concentrations of spores in the suspension corresponded to different treatments. A control treatment contained no spores. Each treatment was applied to a single tray of plants that contained four leaf sets with reference membranes placed on both sides of each leaf set. The number of leaves per treatment ranged from 22 to 30. After inoculation, plants were enclosed in plastic bags for three days to keep them at a high humidity. Plants were maintained in the greenhouse until the time when observation of new lesions was no longer possible due to coalescence of lesions or the onset of natural senescence.

After inoculation, reference membranes were transferred from infection plates to yeast-malt-agar (12 g/l bacteriological agar, 4 g/l yeast extract, 4 g/l malt extract, 4 g/l sucrose, 50 mg/l kanamycin) and incubated for 5-6 days in darkness at 18° C. After incubation, we measured the density of *Z. tritici* colonies by counting them within two or three squares of 1 cm^2^ on digital images of each reference membrane (Appendix S1, Fig. S1 shows examples of colony images). In the second experiment, in treatment with 10^5^ spores/ml we used 0.25 cm^2^ squares. To determine the nature of units that formed fungal colonies, we inspected them under the microscope. When we sprayed the same spore suspension on a microscopy slide during the inoculation, we observed only individual spores without any clumps (we observed 82 spores in total, Appendix S1, Fig. S2). This indicated that each colony grew from an individual spore and therefore individual spores acted as colony forming units (CFUs). At the same time, colony densities did not differ significantly between reference membranes that belonged to the same treatment indicating that the density of spores was the same on reference membranes and on the leaves in their vicinity. We used the colony density measured in this way as an estimate of the density of viable spores in the inoculated areas of leaves assuming that each viable spore was able to form a colony on rich medium under favourable conditions. We provide more details on the inoculation method in Appendix S1.

### Observation of leaves and lesions

To estimate the number of CFUs that landed on each leaf, the areas of leaves were measured. Photographs of leaf sets were taken on the day of inoculation and leaf areas were then measured using Adobe Photoshop CS6. Numbers of CFUs on each leaf were calculated by multiplying the mean densities of colonies on reference membranes on each side of the leaf set by the areas of individual leaves.

To observe the development of lesions, we inspected the infected leaves every day. After the onset of lesion appearance, digital images of leaf sets were captured every day until the majority of leaves became covered by lesions or naturally senescent. Lesions were counted manually on digital images. Newly appeared lesions were marked in the images on the day of their first appearance and the markers were transferred to images captured on the following days in order to avoid double counting of lesions. The total number of lesions on each leaf was calculated from the data on daily appearance of lesions. The lesion density (number of lesions per cm^2^ of leaf area) in each treatment was calculated for each leaf separately based on the total number of lesions that appeared until the last day of observation [34 days post inoculation (DPI) for the first experiment and 36 DPI for the second experiment]. Several leaves were damaged during the experiment and removed.

### Data analysis

Analysis of the data was conducted in R (R Core Team, 2017). We evaluated differences within treatments and between treatments in terms of lesion density and spore density (density of viable spores on the leaf surface). For this purpose, we used two separate analyses of variance (ANOVA). To estimate the infection efficiency of spores that were applied to the leaves, we performed linear regression for the dependency between the spore density and the lesion density on the leaf surface. At high concentrations, we expected to observe a saturation of this dependency because there is a limit to the number of lesions that can form on a single leaf due to finite space and/or resources. For this reason, we used the Akaike information criterion (AIC) to compare a linear model that does not include saturation to a nonlinear model that accounts for possible saturation. AIC balances the goodness of fit with the number of parameters (or complexity) of the model. In this way, we determined whether saturation occurred at high spore densities and identified the range of spore densities that is not affected by saturation. As the nonlinear model, we used the classical Michaelis-Menten model, y=ax/(1+bx), where y is the lesion density, x is the spore density, and a and b are the model parameters. As the linear model, we used the function y=ax. We fitted the two models to the data of each experiment with the nonlinear least squares minimization method (routine nls in R). The data points at high densities that were in the range of saturation according to the AIC score were excluded and the infection efficiency was estimated as the slope of the best-fitting linear function.

To compare the infection efficiency between the two experiments, we tested whether the two slopes were significantly different. First, we used the analysis of covariance (ANCOVA) to test whether there was a significant interaction between the slope and the experiment. Second, we tested whether the two slopes were different using the t-test for the slope difference (Zar, 2010).

## Results

### Experimental design

The spore suspension appeared to be uniformly distributed across the surfaces of leaves and reference membranes when spraying with the paint gun sprayer. The uniformity of the inoculum was quantified by measuring the spatial distribution of fungal colonies on references membranes. The colony densities on reference membranes that belonged to the same treatment exhibited a slight variation but did not differ significantly (ANOVA, p-values between 0.087 and 0.92). As expected, the colony density (averaged over the reference membranes belonging to the same treatment) mostly increased monotonically with increasing the concentration of spores in the suspension (inoculum concentration), as can be seen from Table 1. Only the second lowest spore concentration in the second experiment exhibited a non-monotonic pattern. The spraying resulted on average in 1-2 colonies/cm^2^ on the reference membranes per 1000 spores/ml in the spore suspension (Appendix S1, Fig. S3).

The elastic threads kept the leaves sufficiently flat and stable during the progress of the disease, which enabled accurate observation of the inoculated area of leaves over time.

**Table 1.** Density of spores and density lesions on infected leaves measured on the final day of observation. The values represent means over leaves belonging to the same treatment and the uncertainties represent standard errors of means.

| Treatment | First experiment |  | Second experiment |  |
| --- | --- | --- | --- | --- |
|  | spores/cm<sup>2</sup> | lesions/cm<sup>2</sup> | spores/cm<sup>2</sup> | lesions/cm<sup>2</sup> |
| 5x10<sup>2</sup> | 1.30 ± 0.26 | 0.0922 ± 0.021 | 6.53 ± 0.57 | 0.245 ± 0.036 |
| 10<sup>3</sup> | 2.40 ± 0.31 | 0.101 ± 0.025 | 4.13 ± 0.38 | 0.0660 ± 0.015 |
| 5x10<sup>3</sup> | 7.50 ± 0.58 | 0.368 ± 0.036 | 14.8 ± 0.93 | 0.467 ± 0.049 |
| 10<sup>4</sup> | 21.1 ± 1.0 | 0.742 ± 0.059 | 32.3 ± 2.7 | 1.04 ± 0.090 |
| 5x10<sup>4</sup> | 53.2 ± 1.2 | 2.06 ± 0.056 | 85.9 ± 6.8 | 3.84 ± 0.10 |
| 10<sup>5</sup> | 117 ± 1.7 | 2.75 ± 0.11 | 223 ± 6.8 | 5.24 ± 0.14 |

### Appearance of lesions

According to our observations, lesions were continuously appearing on leaves during the time span of about three weeks (Fig. 2). Figures 2a and 2b show the rate of appearance of new lesions over time, while Figures 2c and 2d show the change over time in the total number of lesions. In every treatment, the dynamics was qualitatively similar. First, lesions started to appear at a slow rate, then the rate of their appearance increased, reached its maximum and eventually dropped to zero (Fig. 2a, 2b). After this time, no new lesions appeared, hence the total number of lesions remained constant (Fig. 2c, 2d).

At higher spore densities, lesions started to appear earlier compared to lower spore densities [compare for example purple curves (squares) to cyan curves (stars) in Fig. 2; the difference was not tested statistically]. Lesions that appeared earlier started to grow and merge with lesions that appeared in their vicinity a few days later (Fig. 1). Without daily observations, we would not be able to distinguish them. Leaves inoculated with higher concentrations of spores carried larger final numbers of lesions. For this reason, in treatments with highest spore densities, lesions covered the entire leaf surface soon after their appearance and we could not observe any further appearance of lesions (Fig. 2). On the contrary, in treatments with lower spore densities, numbers of lesions were smaller and we could observe individual lesions over a longer time.

**Figure 2.**
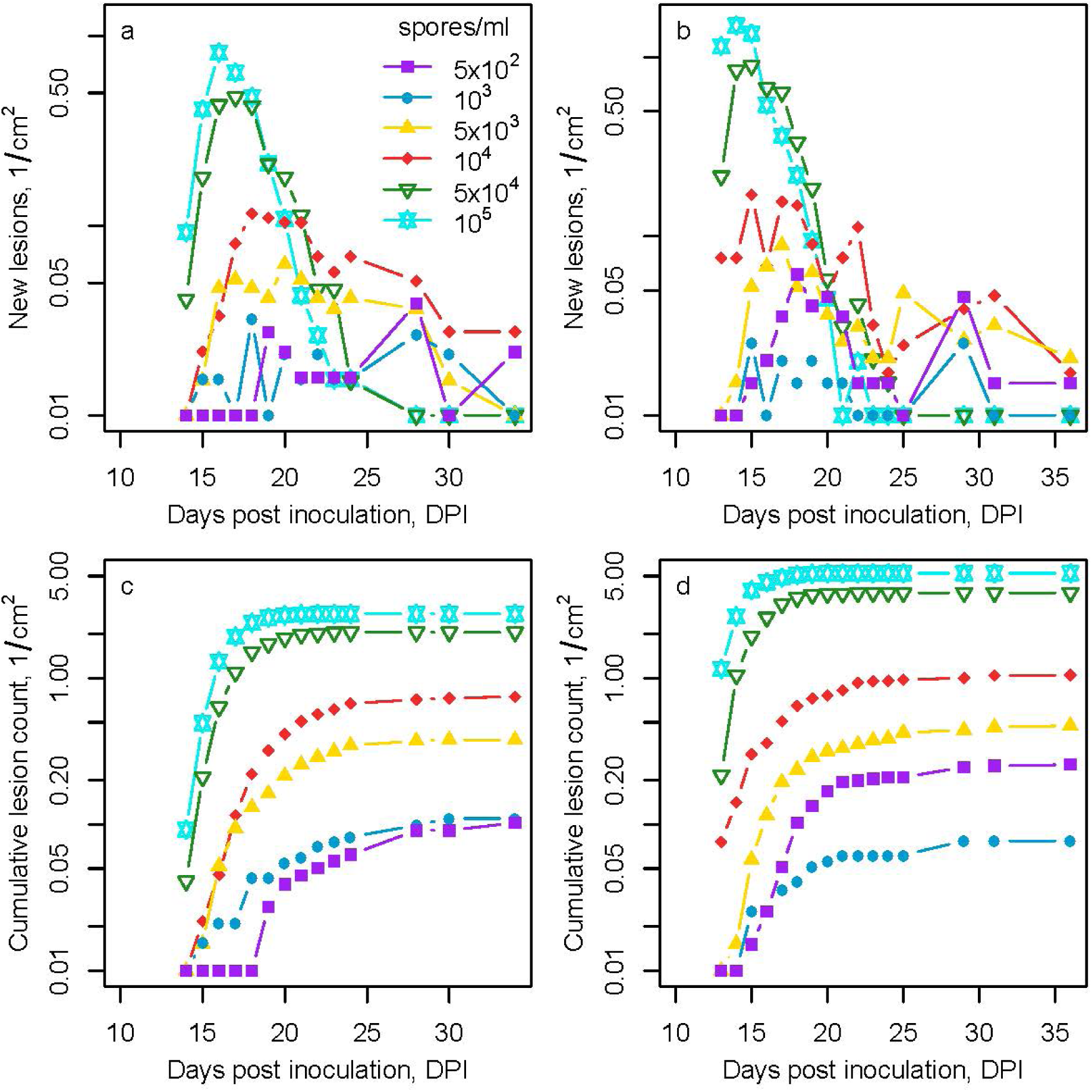
Appearance of lesions. The number of lesions that appeared since the previous observation as a function of time in the first experiment (a) and in the second experiment (b). Total number of lesions in the first experiment (c) and in the second experiment (d). The values were divided by the total leaf area separately in each treatment. Note the logarithmic scale on the y-axis; values are added by 0.01 to show zeros. Different colors/symbols represent treatments with different spore concentrations in the inoculum suspension: purple squares (5×10^2^ spores/ml); blue diamonds (10^3^ spores/ml); yellow filled triangles (5×10^3^ spores/ml); red diamonds (10^4^ spores/ml); green open triangles (5×10^4^ spores/ml); cyan stars (10^5^ spores/ml).

### Measurement of infection efficiency

To determine the infection efficiency, we characterized the numbers of viable spores applied to the leaves and measured the numbers of lesions that subsequently appeared on the leaves (Table 1). In different treatments, the mean spore density (the number of spores per unit leaf area) ranged from 1.3 to 117 spores/cm^2^ in the first experiment and from 3.4 to 223 spores/cm^2^ in the second experiment. The mean lesion density (the number of lesions per unit leaf area) measured on the final day of observation also varied between treatments. This quantity ranged from 0.092 to 2.8 lesions/cm^2^ in the first experiment (measured at 34 DPI) and from 0.066 to 5.2 lesions/cm^2^ in the second experiment (measured at 36 DPI).

Comparison between linear and nonlinear models based on AIC favoured the nonlinear dependency of the lesion density on the spore density. The AIC scores were 223 and 108 in the first experiment, 446 and 300 in the second experiment for the linear and the nonlinear models, respectively. This means that the dependency exhibits a substantial saturation (dashed lines in Fig. 3a). To determine the range of spore densities that is not affected by the saturation, we excluded the treatment with the highest spore densities (117—119 spores/cm^2^ in the first experiment and 212—242 spores/cm^2^ in the second experiment). Now, the comparison favoured the linear model (AIC score −31.1 and −29.8 in the first experiment; 157 and 159 in the second experiment for linear and nonlinear models, respectively). This indicates that saturation only played a role at the highest spore densities, while at lower spore densities the lesion density depended linearly on the spore density and was not affected by saturation. We used this range (all data except for the treatment with 10^5^ spores/ml that corresponds to highest spore densities) to determine the infection efficiency.

We estimated the infection efficiency as the slope of the linear part of the relationship between the lesion density and the spore density (solid lines in Fig. 3). This yielded the estimates (3.8 ± 0.1) % in the first experiment (R^2^ = 0.95) and (4.3 ± 0.2) % in the second experiment (R^2^ = 0.95), where the uncertainties represent 95 % confidence intervals. According to ANCOVA, the slopes differed between the two experiments (p = 1.1 × 10^−6^, F = 24.7, df = 1) and the t-test (Zar, 2010) showed the same result (p = 1.8 × 10^−4^, t = 4.6, df = 327). The full dataset can be accessed from the Dryad Digital Repository xxx.

**Figure 3.**
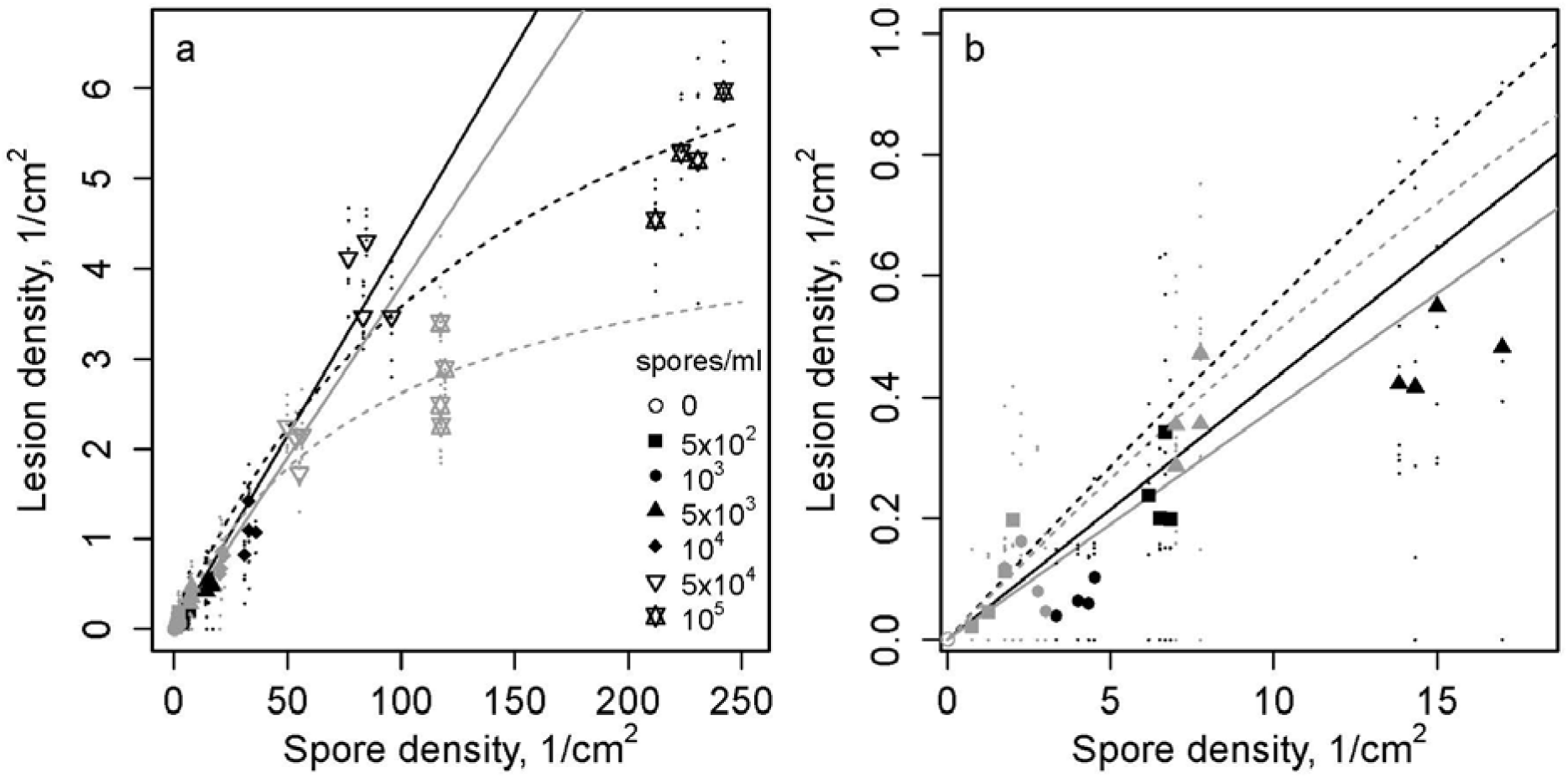
Infection efficiency of *Z. tritici* spores. (a) The lesion density is plotted against the spore density; (b) magnified view of the low-density part of panel (a). Data and curves are shown in grey for the first experiment and in black for the second experiment. Different symbols correspond to treatments with different concentrations of spores in the inoculum, same as in Fig. 2. Dashed lines show the nonlinear function y=ax/(1+bx) fitted to full data of each experiment. Solid lines show the linear function y=ax fitted to data below 100 spores/cm^2^.

## Discussion

We have presented an accurate and reliable method for measuring the infection efficiency of *Z. tritici*. The main advantages of this method include the use of reference membranes to estimate densities of viable spores deposited on leaves, and daily imaging of leaves attached to metal plates. Counting colonies on reference membranes allowed us to estimate densities of inoculated, viable spores accurately and confirmed the uniformity of their spatial distribution on leaf surfaces. Attaching leaves to metal plates allowed for efficient daily observations that resulted in accurate counting of individual lesions. The resultant stability of the leaves also ensured that the inoculum did not move across the leaf surface, and facilitated the observation of new lesions in daily photographs. The method is based on commonly used, low-cost equipment, which makes it affordable and easy to modify.

Each of the individual components of our method has been used in the literature previously. Similar ideas concerning the estimation of viable spore densities with reference plates were used by Chakraborty *et al*. (1990) in a different pathosystem; a range of inoculum concentrations, including low concentrations, were used by Fones *et al*. (2015) in this pathosystem; attached leaf assays were previously used in this pathosystem for example by Keon *et al*. (2007) and Lee *et al*. (2014); time-resolved imaging was used before in fungal biology for example in studies of spore ejection of basidiomycete fungi (Noblin *et al*., 2009). In addition, numerous studies measured the ability of fungal spores to grow colonies under conducive conditions *in vitro* to estimate the viability of spores: for example Valsecchi *et al*. (2017) in *Aspergillus fumigatus* and King *et al*. (2017) in *Z. tritici*. Unlike the previous studies in this pathosystem that used cotton swabs to deploy the inoculum suspension on leaves (Lee *et al*., 2014) or pipetted a well-defined volume of spore suspension onto the leaf and spread it using a gloved finger (Fones *et al*., 2015), we used a paint gun sprayer [comparable to Statler & McVey (1987) for example] that produced an evenly distributed “fog” of tiny droplets. This allowed us to characterize the density of viable spores and to verify the uniformity of their spatial distribution using reference membranes. Thus, the novelty of our work lies not so much in each individual component of the method *per se*, but rather in a specific way we combined these methodological aspects that allowed us to obtain accurate estimates of the pathogen’s infection efficiency.

In contrast to some of the earlier measurements of infection efficiency in plant pathogens that quantified the *total* number of spores in the applied inoculum, we estimated the number of *viable* spores in the applied inoculum. This is because when estimating the infection efficiency, we need to quantify the proportion of *viable* spores that have caused lesions. We estimated the density of viable spores from the density of fungal colonies on agar plates (as described in Materials and Methods). Depositing spores of fungal pathogens on solid medium, incubating them under optimal conditions and counting the number of resulting colonies is an established method to evaluate the viability of fungal spores, which has been previously used in Z*. tritici* (King *et al*., 2017) and in other pathogens, for example in *Aspergillus fumigatus* (Valsecchi *et al*., 2017). Viability of spores is likely to be higher on agar than on the leaf surface because (i) agar plates contain more nutrients and (ii) plates are maintained under constant environmental conditions that are close to optimal for the fungus in contrast to the leaf surface in the greenhouse that is exposed to various stress factors such as variable humidity, temperature and light with a wide frequency spectrum including UV. This difference is acceptable because our goal was to estimate the maximum number of viable spores in the sprayed suspension. Decreased survival of spores on leaves compared to agar is included in the estimate of the infection efficiency. At the same time, there could be factors that promote germination and thus the observed viability of the spores on the leaf surface. However, to our knowledge, there is no evidence for host-specific cues or other factors promoting germination of *Z. tritici* spores on the leaves. For this reason, we assumed that the maximum viability of spores was captured by the method described.

As described above, we quantified the infection efficiency as the proportion of *in vitro* viable spores that have caused lesions. However, infection efficiency can be defined in several different ways and the choice of the definition should be determined by the aims of the study. We can think of four definitions listed from the most inclusive to the most exclusive: (i) the proportion of all spores that have caused lesions, (ii) the proportion of *in vitro* viable spores that have caused lesions, (iii) the proportion of spores germinated on the leaf (i.e., spores viable *in planta*) that have caused lesions, (iv) the proportion of penetrating spores that have caused lesions. The difference between (i) and (ii) results from dead spores present in the inoculum. The difference between (ii) and (iii) arises because the definition (ii) takes into account environmental factors that may cause differences in viability between optimal conditions *in vitro* and likely more harsh conditions *in planta*. Therefore, measuring the difference between (ii) and (iii) could be used to quantify abiotic factors affecting infection efficiency. In contrast to (iii), the definition (iv) excludes the penetration efficiency of the spores and measures the infection efficiency of the spores that have penetrated successfully. Using definitions (iii) and (iv) would lead to a more precise quantification of the host-pathogen interaction compared to definitions (i) and (ii) as this would exclude some of the purely environmental variability and disentangle different components of infection success (Fones *et al*., 2015). Measuring infection efficiency according to definitions (iii) and (iv) is more difficult than using the definition (ii), but it could be done e.g. with the help of methods presented by Fones *et al*. (2017). Such measurements would also provide more detailed information on the component of plant’s quantitative resistance that decreases the infection efficiency of *Z. tritici*. However, the definition (ii) better corresponded to the aims of our study for two reasons. First, the mortality of spores in the inoculum suspension can be influenced by a range of factors not related to host-pathogen interaction. Hence, we think that by excluding dead spores we achieved a more reliable measure of the interaction between the host and the pathogen compared to using the definition (i). Second, in contrast to definitions (iii) and (iv), the definition (ii) includes environmental factors responsible for the difference in spore viability between *in vitro* and *in planta* conditions, which are relevant for the epidemiology of the pathogen.

We found that the relationship between the density of fungal colonies on reference membranes and the concentration of spores in the suspension was monotonic: colony density increased at higher concentrations of spores. An exception to this tendency was observed only for the second lowest spore concentration in the second experiment, which exhibited a non-monotonic pattern (see Table 1). This non-monotonic pattern could result from a non-homogeneous distribution of spores in the suspension during the spray, if the spores had time to settle down inside the sprayer tank. At low concentrations of spores, the dependency may also be sensitive to slight variations in the manual movement of the sprayer between treatments. Noticeable deviations from a perfect linear fit between the spore concentration in the suspension and the density of colonies on plates were observed for example in treatment 10^4^ spores/ml, which resulted in higher colony density than expected (Appendix S1, Fig. S1). All these observations indicate that the measurement of the concentration of spores in the inoculum suspension is not sufficient to determine reliably the actual density of spores on leaf surfaces. Therefore, an important aspect of the method is to characterize the density of viable spores present on the leaves after every spray of the inoculum suspension. We achieved this conveniently by counting colonies on reference membranes.

Our measurements show that lesions did not all appear within a narrow time span of a few days. On the contrary, they were continuously appearing during a period of up to three weeks (see Fig. 2), even though we used a single pathogen strain to infect a single host variety. This observation is consistent with recent modelling of distributions of incubation periods (Ottino-Loffler *et al*., 2017), but it contradicts the established view in the literature on *Z. tritici* according to which lesions appear after an asymptomatic period of approximately two weeks (Kema *et al*., 1996; Steinberg, 2015). Shaw (1990) used pycnidiospores from the natural field population of *Z. tritici* to inoculate wheat plants in the greenhouse and found that lesions were appearing continuously during the period of up to 25-30 days. However, in the experiments of Shaw (1990), the asynchronicity in lesion appearance could result from both the variation in the latent period between different pathogen strains and the “developmental” asynchronicity within a single pathogen genotype that we observed here. Recently Fones *et al*. (2017) have shown that there is a large variation in the timing of penetration events of spores originating from the same inoculation event. They observed spores on leaf surfaces until 10 DPI and found that the fraction of successfully penetrating individuals was continuously increasing. This variation in the duration of the epiphytic growth offers a possible explanation for large differences in the times of appearance of individual lesions that we observed. At higher inoculum concentrations, lesions started to appear earlier possibly due to a higher total number of infections that led to an increase in the number of rare “fast” infections. Consequently, the large variation in the duration of the asymptomatic period that we observed at low inoculum concentrations is likely hidden in experiments that use high inoculum concentrations [e.g., 10^6^ spores/ml is typically used in greenhouse trials, for example in (Stewart & McDonald, 2014)]. In this case, leaves become fully necrotic only a few days after the appearance of first lesions and the appearance of new lesions is no longer visible.

In treatments with low densities of only 1-5 spores/cm^2^, it is highly plausible that individual lesions were caused by single spores. This is consistent with experimental findings of Shaw (1990), in which the inoculations were conducted with the population of natural field strains of *Z. tritici*. Fones *et al*. (2015) confirmed the ‘one spore, one lesion’ hypothesis using the reference strain of *Z. tritici* IPO323. This may not be the case at higher spore densities when the area covered by a typical lesion contains many viable spores and the spores may cooperate to cause infection and form a lesion. Indeed, *Z. tritici* is able to undergo anastomosis between germinating spores when they are deposited on the leaf close to each other (Mehrabi *et al*., 2009). Spores may also interact in more subtle ways, as was observed in other foliar plant pathogens (Jeffries, 1995, Jesus Junior *et al*., 2014). For example, when two spores penetrate the leaf surface at nearby locations, the probability of lesion formation may become higher (cooperation) or lower (antagonism) than twice the probability to form a lesion by an individual spore that is far away from other spores. Also, when one spore causes a lesion, the probability of other spores in the vicinity to cause a lesion may increase (cooperation) or decrease (antagonism). Such interactions should lead to departures from a linear dependency between the density of spores and the density of lesions of leaves. According to our measurements, this dependency is linear within a wide range of spore densities, from low densities of 1-5 spores/cm^2^ to intermediate densities of 10-80 spores/cm^2^. This indicates that in this range of spore densities, lesions are caused by single spores with no evidence of interaction between spores within the limits of accuracy of our measurements. Interaction between spores may still occur at higher spore densities, but in our experiment, its effect would not be visible because of the saturation of the leaf surface with lesions.

Some evidence for a possible cooperation between spores belonging to the same strain in causing lesions was provided by Haueisen *et al*. (2017). After inoculating wheat leaves with a suspension containing a high concentration of *Z. tritici* spores (10^8^ spores/ml), they found cases when several hyphae entered a single stoma. Additionally, noticed that the spatial distribution of stomatal penetrations appeared to be clustered, rather than uniform. Some indication for an antagonistic interaction between different *Z. tritici* strains in pycnidia formation was reported by Schuster *et al*. (2015): each pycnidium was produced by a single strain when two strains were inoculated together even though hyphae of the two strains were found growing next to each other.

Our estimates of infection efficiency were close to 4 % in both biological replicates. However, statistical tests revealed that the difference in the estimates between the two replicates was significant. Several factors may be responsible for the difference. The experiment was conducted under controlled greenhouse conditions but changes in the external weather may still have affected the outcomes. While the first experiment was performed in early January, the second experiment was performed in the beginning of March. Hence, both the amount of external light and the external temperature differed between the experiments. In addition, during the second experiment, we observed several times water droplets on infected leaves and aluminium plates. This may have caused microclimatic differences between the two replicates and such differences are known to affect infection success of *Z. tritici* (Shaw, 1990). The fungal inoculum was grown from the same batch and prepared with the same protocol in both replicates, but the fungus is capable of rapidly accumulating genetic changes through mitotic events (Möller *et al*., 2018) and this may have contributed to the difference in the estimates of infection efficiency.

Infection efficiency of *Z. tritici* blastospores of strain IPO323 was estimated to be around 50 % on wheat cultivar Galaxie by Fones *et al*. (2015), which is much higher than the estimate of around 4 % that we report here. Many factors may have contributed to this difference. Aggressiveness of the two pathogen strains may be different as well as the degree of resistance of the two wheat cultivars. Greenhouse conditions in the experiment by Fones *et al*. (2015) differed from our experiment in terms of light and temperature. Also, the age of infected seedlings was only 10 days compared to 16 days in our experiment. Another major factor could be the difference in the inoculation methods. They pipetted droplets of the spore suspension on the leaf surface and spread them with a gloved finger, while we sprayed the spore suspension creating a fog consisting of microscopic water droplets. When spread with a finger, spores were likely to be placed close to the leaf surface. In contrast, a substantial fraction of water droplets in the fog may have remained on trichomes (leaf hairs), which could diminish their penetration success. Our estimate for infection efficiency of *Z. tritici*, about 4 %, lies within the lower range of what was previously reported in the literature for other fungal pathogens. Among 22 infection efficiency estimates in different species discussed by Sache & de Vallavieille-Pope (1995), eight species had the values within the range of 0–5%, five species ranged within 6–15%, five species ranged within 16–25% and four species ranged within 26–50%. Notably, Sache & de Vallavieille-Pope (1995) used the highest value for each species they found in the literature. More recent research on measuring the infection efficiency of fungal plant pathogens is scarce, but Li *et al*. (2010) estimated the infection efficiency of soybean rust (*Phakopsora pachyrhizi*) to be within the range of 0.5–10%.

We infected a single wheat variety with blastospores of a single strain of *Z. tritici* to provide a proof of principle for the reliable measurement of the infection efficiency. Using blastospores is a standard method to conduct greenhouse trials with this pathogen, but the role of blastospores in the pathogen’s life cycle remains unknown. However, our method could also be used to measure the infection efficiency of both pycnidiospores and ascospores, which are known to drive the epidemics in the field. Infection efficiency may be different in other pathogen strains or when infecting other wheat varieties. Haueisen *et al*. (2017) observed that three strains of *Z. tritici* differed in terms of the time between the inoculation and the first stomatal penetration and in terms of the degree of epiphyllous proliferation. We also expect to observe specialization of pathogen strains to certain wheat varieties, because the pathogen population is extremely diverse (Linde *et al*., 2002) and is known to adapt rapidly to different host environments (Poppe *et al*., 2015). These factors may lead to different infection efficiencies in different *Z. tritici* strains. Therefore, we do not expect that our estimate of infection efficiency in this particular pathogen strain-host cultivar combination would be representative of the natural pathogen population. However, using this method, a number of pathogen strains can be tested on a number of wheat varieties. Such a comprehensive characterization of infection efficiency would improve the predictive power of mathematical models that describe epidemic development and pathogen evolution and in this way contribute to improving control strategies. If a sufficient degree of heritable variation in terms of infection efficiency is found in the pathogen population, the genetic basis of this trait could be revealed, for example by conducting a genome-wide association study. Similarly, from the perspective of the host, our method opens opportunities for determining the genetic basis of the component of quantitative resistance that suppresses infection efficiency. This knowledge has the potential to improve and accelerate breeding for quantitative resistance against STB.

## Acknowledgements

P. Karisto and A. Mikaberidze gratefully acknowledge financial support from the Swiss National Science Foundation through the Ambizione grant PZ00P3_161453. The authors would like to thank Bruce A. McDonald and the members of the Plant Pathology group at the ETH Zürich for fruitful discussions.

## Appendix S1. Spraying results

Figure S1 shows examples of digital images of reference membranes containing fungal colonies of *Z. tritici*. Since the individual colonies can be clearly distinguished from these images, by counting the number of colonies per unit area we were able to determine both the density and the spatial distribution of the colony forming units (CFUs) across the inoculated area.

Figure S2 shows microscopic images of spores on a microscopic slide after inoculation. We see that spores do not form clumps even though they are sometimes located close to each other. At lower spore concentrations average distance between individual spores is expected to increase (not tested by microscopy). This supports our assumption that colonies on reference membranes were formed by individual spores. We counted in total 82 spores on the microscopy slides. None of the spores were clearly in contact to other spores, although 17 of them showed signs of budding and would likely have produced new spores later if incubated further. Budding of the blastospores is an inherent property of spore culture grown under conducive conditions and it is the way they multiply *in vitro*.

Figure S3 depicts how the density of colonies on reference membranes depends on the concentration of spores in the inoculated suspension. The dependency shows a monotonic increase in both experiments, with the exception of the two lowest spore concentrations in the second experiment, which exhibited a non-monotonic pattern (see Table 1 in the main text). The occasional deviations from linear fit can result from differences in the action of spraying. These variations highlight the importance of the reference membranes.

**Figure S1.**
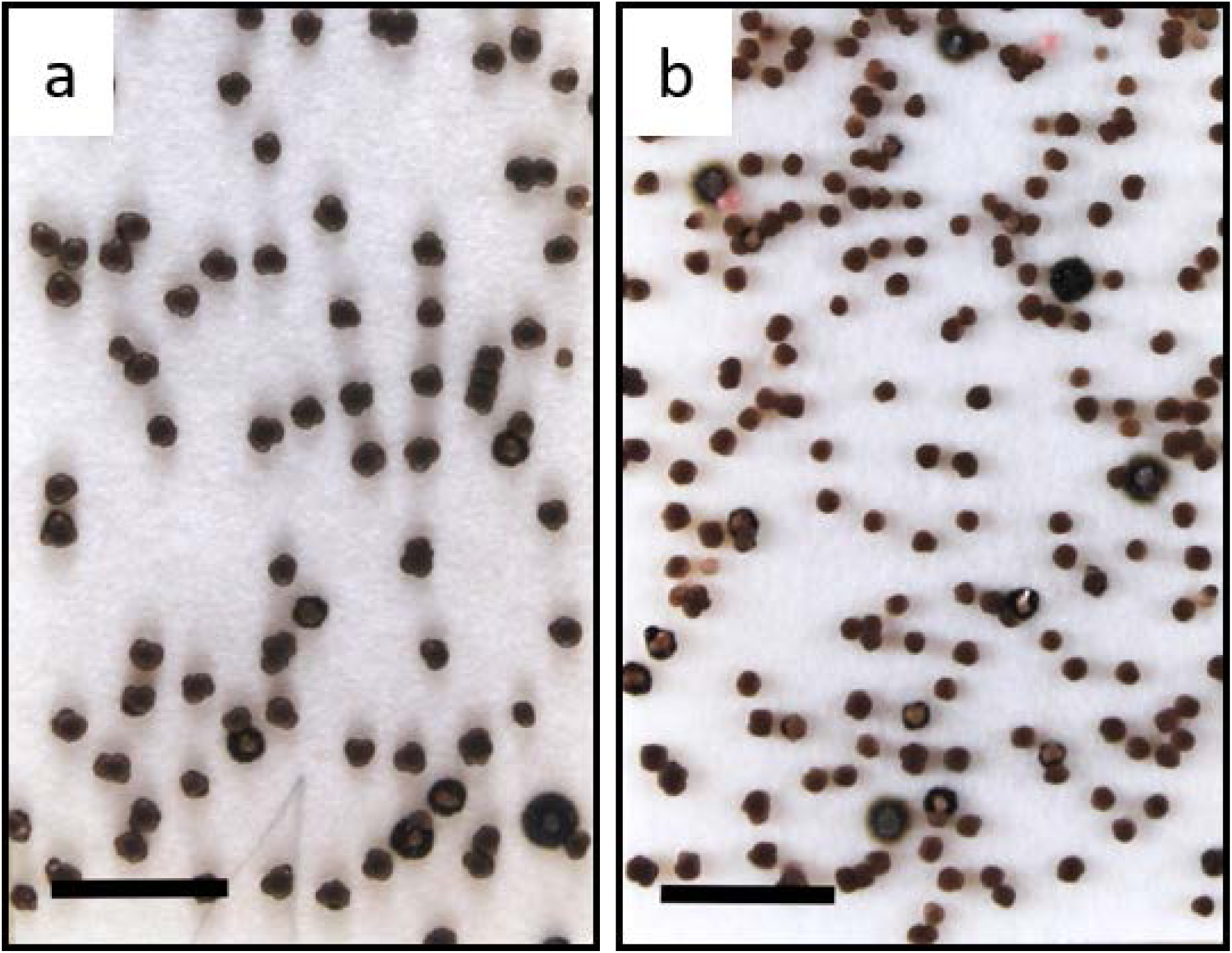
Representative reference membranes showing the density and the spatial distribution of *Z. tritici* spores applied to the leaves. (a) Treatment with 5 × 10^2^ spores/ml; (b) treatment with 5 × 10^3^ spores/ml. Size of scale bar is 1 cm, photographs were taken at a somewhat later time point than the time when the colonies were counted.

**Figure S2.**
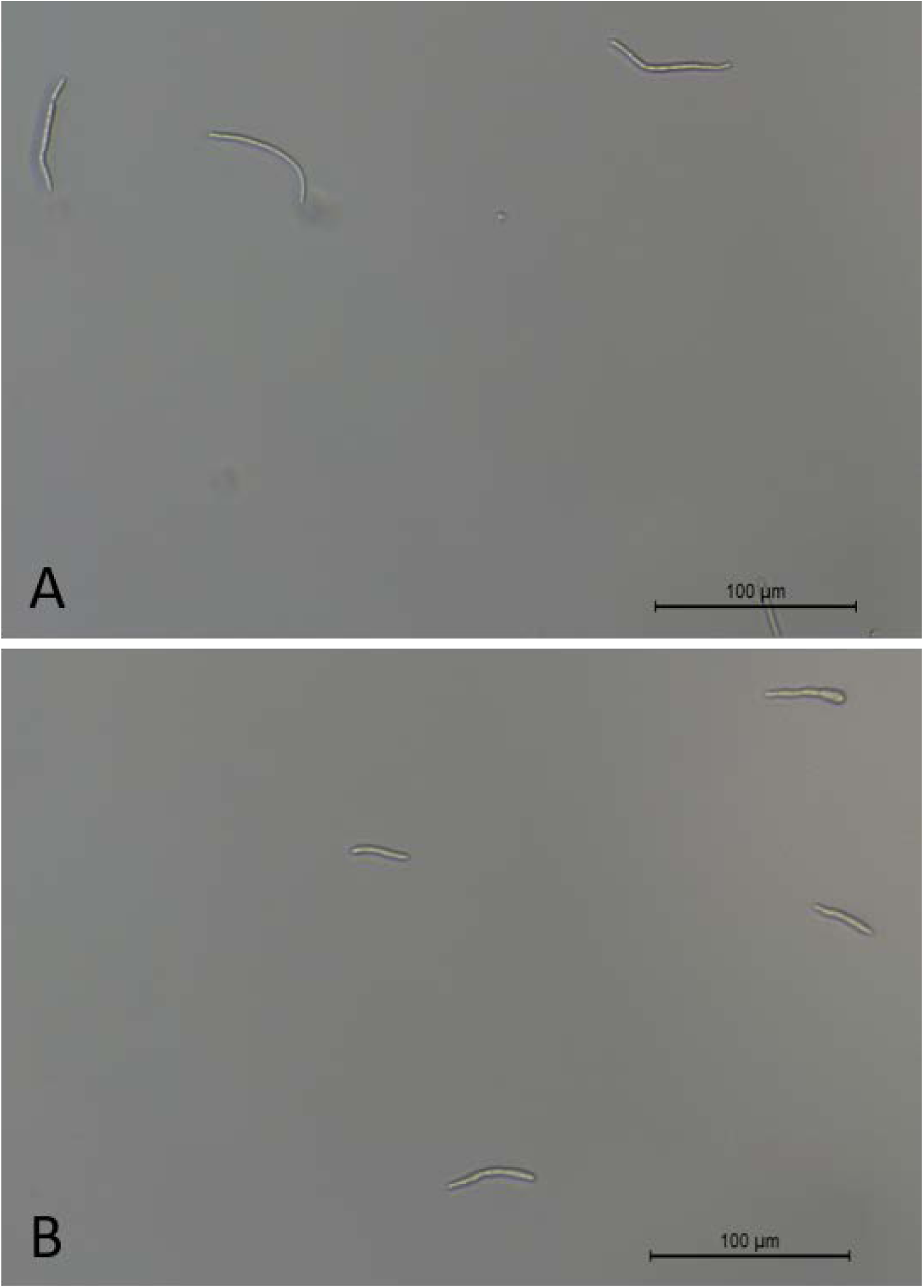
Microscopic images of spores after spraying the 10^5^ spores/ml suspension to a microscopic slide. Panels (a) and (b) show different sections of the microscopic slide.

**Figure S3.**
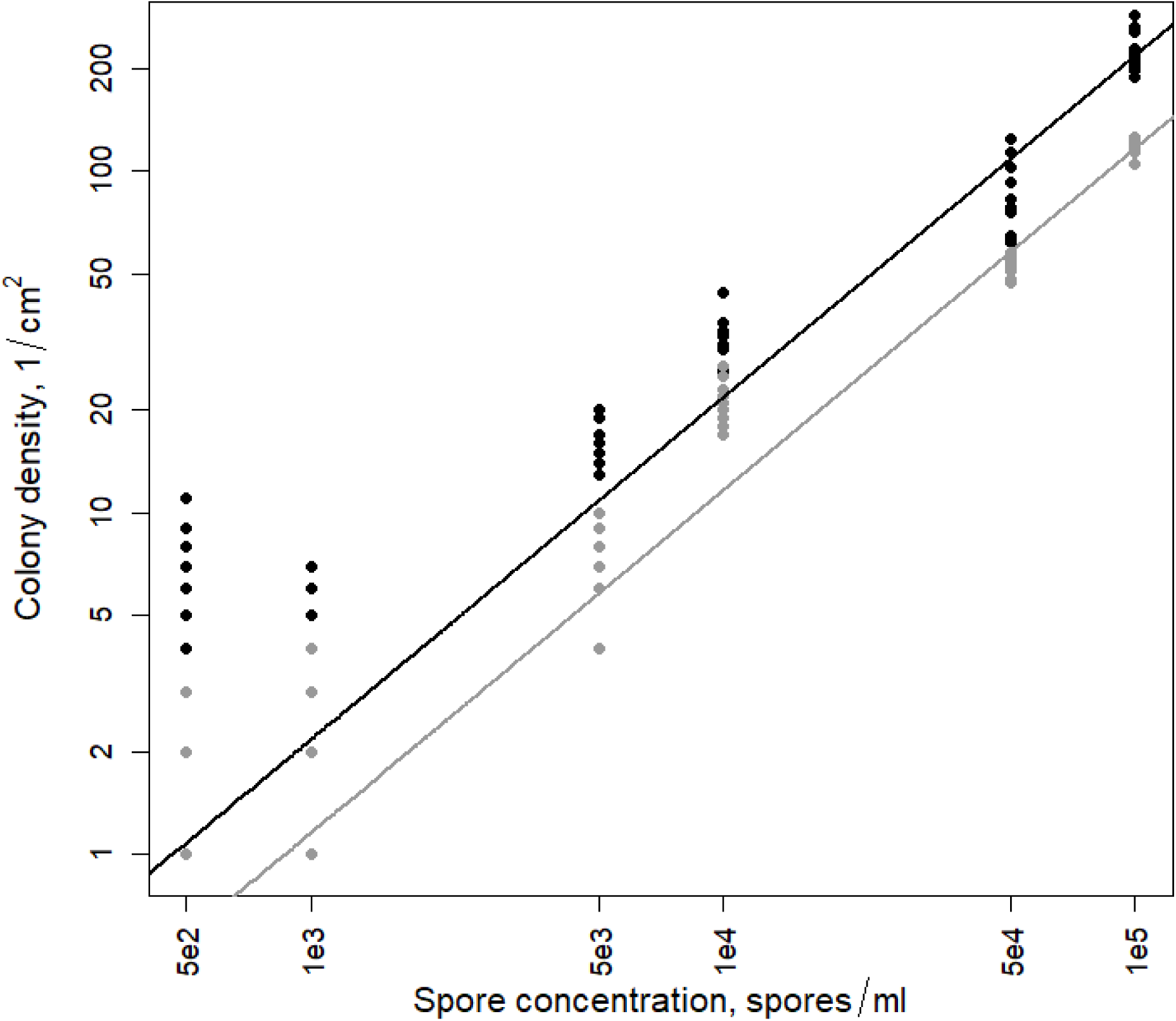
Dependency of the colony density on the spore concentration in the inoculum suspension in the first experiment (grey) and in the second experiment (black). Data points resulting from spore counts within individual squares on reference membranes are shown with circles. Lines correspond to best fits of linear functions with zero intercepts. The fits were conducted on the linear scale. The slopes are 0.0012 (colony/cm^2^)/(spore/ml) (p < 2 × 10^−16^, R^2^ = 0.99) and 0.0022 (colony/cm^2^)/(spore/ml) (p < 2.2 × 10^−16^, R^2^ = 0.97) for the first and second experiment respectively. Note the logarithmic scale on both axes.

